# Experimental validation of computational models for the prediction of phase distribution during multi-channel transcranial alternating current stimulation

**DOI:** 10.1101/2023.04.07.536090

**Authors:** Sangjun Lee, Sina Shirinpour, Ivan Alekseichuk, Nipun Perera, Gary Linn, Charles E. Schroeder, Arnaud Y. Falchier, Alexander Opitz

## Abstract

Transcranial alternating current stimulation (tACS) is a widely used noninvasive brain stimulation (NIBS) technique to affect neural activity. Neural oscillations exhibit phase-dependent associations with cognitive functions, and tools to manipulate local oscillatory phases can affect communication across remote brain regions. A recent study demonstrated that multi-channel tACS can generate electric fields with a phase gradient or traveling waves in the brain. Computational simulations using phasor algebra can predict the phase distribution inside the brain and aid in informing parameters in tACS experiments. However, experimental validation of computational models for multi-phase tACS is still lacking. Here, we develop such a framework for phasor simulation and evaluate its accuracy using *in vivo* recordings in nonhuman primates. We extract the phase and amplitude of electric fields from intracranial recordings in two monkeys during multi-channel tACS and compare them to those calculated by phasor analysis using finite element models. Our findings demonstrate that simulated phases correspond well to measured phases (r = 0.9). Further, we systematically evaluated the impact of accurate electrode placement on modeling and data agreement. Finally, our framework can predict the amplitude distribution in measurements given calibrated tissues’ conductivity. Our validated general framework for simulating multi-phase, multi-electrode tACS provides a streamlined tool for principled planning of multi-channel tACS experiments.

## 1. Introduction

Transcranial alternating current stimulation (tACS) is a noninvasive brain stimulation technique (NIBS) that aims to modulate brain oscillations in a frequency-specific manner by applying a weak external current via electrodes attached to the scalp (Wischnewski et al., 2022). Several studies have shown that tACS can improve various brain functions. However, researchers are still investigating the relationship between specific tACS parameters and behavioral outcomes. For instance, in healthy participants, theta tACS improved memory performance (Alekseichuk et al., 2016; Alekseichuk et al., 2020). Moreover, tACS can enhance motor-related excitability (Schilberg et al., 2018), long-term memory consolidation (Ketz et al., 2018), and visual perception (Salamanca-Giron et al., 2021) by adjusting the stimulation frequency corresponding to the targeted brain function. Clinical trials have also shown promising results for tACS. It has been used to treat neurological and psychiatric symptoms such as depression, epilepsy, and schizophrenia (Ahn et al., 2019; Cappon et al., 2016; Haller et al., 2020), stroke rehabilitation (Naros and Gharabaghi, 2017), and Parkinson’s disease (Guerra et al., 2022).

TACS can entrain brain rhythms in a phase-specific manner by synchronizing intrinsic neural oscillations to its applied stimulation phase (Johnson et al., 2020; Krause et al., 2019). Traditionally, tACS is applied with only 0° or 180° phase differences between stimulation electrodes (Polanía et al., 2012). This results in a standing wave electric field, which synchronizes brain areas with zero phase lag. The phase dependent large-scale synchronization of brain oscillations across distinct brain regions plays a crucial role in brain function (Fries, 2015). Thus, standing wave tACS can be used to manipulate the phase alignment between distinct regions to strengthen or weaken a functional connection in the brain. This can be accomplished by injecting two alternating currents with a phase difference of either 0° or 180°. Previous studies demonstrated that 0° (in-phase) tACS over the frontal and parietal cortex for targeting the frontoparietal network improved working memory performance (Draaisma et al., 2022; Violante et al., 2017). On the contrary, 180° (anti-phase) tACS over the same regions deteriorated a cognitive process (Takeuchi et al., 2021).

Using tACS input currents with defined phase shifts results in a traveling wave electric field. Electrophysiological traveling waves refer to time-lag neural oscillation patterns, characterized by a gradual phase shift of neural oscillations across the brain (Bolt et al., 2022; Zhang et al., 2018). Previous studies have shown that brain oscillations in low frequency bands propagate across the cortex in the form of traveling waves, especially during cognition (Halgren et al., 2019; Muller et al., 2018). It has been suggested that traveling waves are a key mechanism for explaining the transfer of information across cortical regions (Muller et al., 2018). Alekseichuk et al. (2019a) showed that multi-channel tACS can produce electric fields in the form of traveling waves inside the brain. They suggested the possibility of traveling wave tACS (twtACS) that can entrain time-lag brain oscillations across remote brain regions.

To predict and optimize the phasic electric field distribution during twtACS, it is necessary to develop and validate computational models. Computational simulations using the finite element method (FEM) have been well established to plan tACS experiments in human studies (Baltus et al., 2018; Klírová et al., 2021; Rufener et al., 2019). These simulation studies have mostly been focused on calculating the amplitude distribution of electric fields during tACS. A previous study demonstrated that the distribution of electric fields varies depending on whether in-phase or anti-phase tACS is used (Saturnino et al., 2017). Our group previously suggested an analytical approach for calculating the electric field phase distribution during multi-channel tACS employing phasor algebra (Alekseichuk et al., 2019a). This allows determining the phase value inside the brain depending on the phase difference of the applied tACS currents. However, a systematic validation and determination of the accuracy of our approach (hereafter referred to as “phasor simulation”) for predicting the phase of electric fields for tACS is still lacking.

Here we compare the phase distribution obtained from phasor simulation and *in vivo* intracranial recordings in two NHPs during multi-channel tACS to validate our computational modeling approach. Two active electrodes were placed on the scalp over the middle forehead and left occipital lobe. The return electrode was placed over the left temporal region. We extracted the phase and the amplitude of electric fields from intracranial recordings in two NHPs under various stimulation phase conditions. Then, we conducted the phasor simulation using head models to calculate phasic electric fields under the same conditions as in recordings, followed by a comparison of the measured and simulated results. In addition, we evaluate the effect of a small displacement of the return electrode, which is essential for the formation of a phase gradient, on the phase distribution in simulations. As FEM simulations have a tendency to overestimate the electric field amplitude (Opitz et al., 2018), we optimize the electrical conductivity of NHP tissues to minimize the error between measured and simulated amplitudes.

## 2. Methods

### 2.1. In vivo experiments in nonhuman primates

#### 2.1.1. Nonhuman primates

All procedures were approved by the Institutional Animal Care and Use Committee of the Nathan Kline Institute for Psychiatric Research. Two NHPs were utilized in experiments. Monkey 1 is a female capuchin monkey (11 years old, and 2.9 kg) whereas monkey 2 is a female rhesus macaque (6 years old, 4.8 kg). In all monkeys, three stereo-EEG (sEEG) (Ad-Tech Medical Instruments Corporation, Racine, Wisconsin, USA) electrodes with 5 mm spacing between electrode contacts were implanted through an entry point in the left occipital cortex. Electrodes were aligned in anterior-posterior direction. One sEEG electrode had an endpoint in the frontal cortex (12 contacts, the other in the medial prefrontal cortex (10 contacts), and another in the anterior hippocampus (10 contacts).

#### 2.1.2. Transcranial alternating current stimulation

tACS protocols and *in vivo* recordings were carried out according to the previous study (Alekseichuk et al., 2019a). tACS using the multi-channel Starstim system (Neuroelectrics, Barcelona, Spain) was applied to the monkeys with two active electrodes placed on the scalp above the forehead (anterior electrode) and over the left occipital lobe (posterior electrode), respectively. The return electrode was attached to the scalp over the temporal region. All the electrodes were round, with a radius of 10 mm. While injecting the alternating current with a fixed phase of 0° from the anterior electrode, we injected the alternating current with the phase varying from 0° to 360° in a step of 15° for the posterior electrode, with the amplitude (peak-to-zero) fixed to 0.1 mA at a frequency of 10 Hz. The current at the return electrode is set so that the sum of all electrode currents always sums to zero. This led to 25 different stimulation conditions with different phase differences between the two active electrodes. For each condition, the stimulation duration and ramping up/down time were 30 s and 5 s, respectively.

#### 2.1.3. Data acquisition and analysis

While injecting tACS, electric potentials in sEEG electrodes were acquired at a sampling rate of 5 kHz using a BrainAmp MR plus amplifier (Brain Products) for monkey 1 and a Cortech NeurOne Tesla amplifier (Cortech Solution, Wilmington, NC) for monkey 2. The raw data were band-pass filtered at cut-off frequencies of 5 Hz and 20 Hz using a 4th-order zero-phase Butterworth filter (Alekseichuk et al., 2019a), then downsampled to 1 kHz using MATLAB 2021b (MathWorks) and the Fieldtrip toolbox (Oostenveld et al., 2011). With the elimination of the 5 s ramping up/down period, 30 s of electric potentials for 25 stimulation conditions were extracted and rescaled to the electric potential when the current of 1 mA was assumed to be injected (Alekseichuk et al., 2019a). Then, we examined preprocessed data and post-implanted MR images to assess for signal contamination and abnormal sEEG contact placement. The contacts outside the gray matter (GM) and white matter (WM) were removed for both monkeys. For monkey 1, three contacts were excluded and interpolated from neighboring contacts. For monkey 2, one sEEG electrode was entirely omitted due to low signal quality. In addition, the front two contacts of the sEEG electrode implanted over the hippocampus were positioned on the boundary between GM and CSF, and substantially twisted relative to the other contacts. Due to the possibility of inaccurate calculations of electric fields resulting from misalignment among contacts, these two contacts were excluded from the analysis. After removing the contacts, a total of 28 contacts from three sEEG electrodes (A1, A2, and A3) and a total of 19 contacts from two sEEG electrodes (B1 and B2) were utilized to calculate the phase (see **Fig. 1A**). Also, three stimulation conditions (0°, 180°, and 360°) were excluded from further analysis because they do not generate phase-gradient electric fields (similar to standard tACS). Note that the aim of this study is to investigate how accurately phasor simulation can predict a tACS-induced phase gradient to manipulate time-lag brain oscillations across brain regions. Thus, 22 stimulation conditions were considered for data analysis. Then, electric fields were calculated using the numerical gradient of measured electric potentials (zero-padded to 2^15^ samples) along contacts for each sEEG electrode. As all sEEG electrodes were aligned in the anterior-posterior direction, we were able to quantify the electric field in the anterior-posterior direction at each contact. Using the Fast Fourier Transform (FFT), we extracted the phase φ and amplitude |***E***| of electric fields at each contact for the maximum frequency (which was equal to the stimulation frequency, 10 Hz) and centered the phase values along each sEEG electrode between –π/2 and π/2 following unwrapping them (Opitz et al., 2016).

**Figure. 1.**
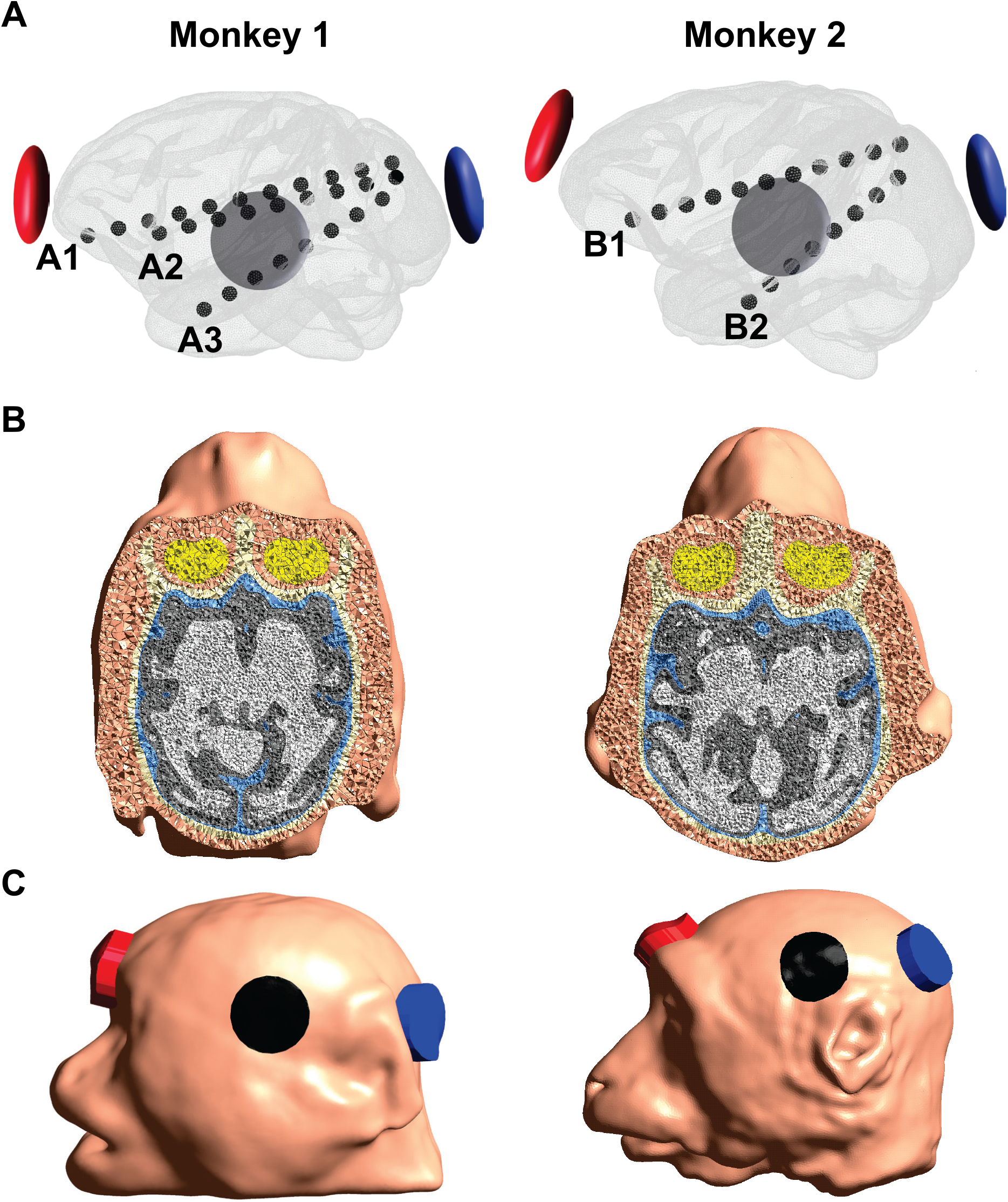
A) Illustration of the location of sEEG electrodes and tACS electrodes. Three and two sEEG electrodes were used for further data analysis for monkey 1 and monkey 2, respectively. B) Illustration of volumetric head models, including the scalp, skull, CSF, GM, WM, and eyes. C) Location of tACS electrodes in head models. The red and blue represent the active electrodes (anterior and posterior electrodes, respectively), and the black represents the return electrode.

### 2.2. Computational simulation for the phase analysis using head models

#### 2.2.1. Realistic finite element model

Realistic head models of the monkey head were created using T1-weighted magnetic resonance (MR) images acquired before sEEG electrodes were implanted. We extracted the GM and WM masks using a modified Human Connectome Project pipeline for non-human primates (Fair et al., 2020). In addition, we performed manual modification for GM and WM masks using ITK-SNAP to include anatomical information that is not captured by automatic segmentation (Yushkevich et al., 2006). The masks for the scalp, skull, cerebrospinal fluid (CSF), and eyes were constructed by manual segmentation. FSL’s FLIRT package was used to register MRI from native space to the Freesurfer space (Jenkinson et al., 2002; Jenkinson and Smith, 2001), and then Gmsh was used to generate a volumetric head model from tissue masks (Geuzaine and Remacle, 2009) (see **Fig. 1B**). We then determined the location of sEEG electrodes used in *in vivo* experiments on head models based on post-implanted MR images for both monkeys.

#### 2.2.2. Phasor simulation

For each monkey model, two active electrodes and the return electrode, with a diameter of 10 mm and a thickness of 5 mm, were attached to the scalp. The location of tACS electrodes was identical to that of *in vivo* experiments. The following conductivity values were used for phasor simulation: 0.465 S/m for the scalp, 0.5 S/m for the eyes, 0.01 S/m for the skull, 1.654 S/m for the CSF, 0.275 S/m for the GM, and 0.126 S/m for the WM. We calculated electric fields inside the brain by solving the Laplace equation given by –∇·(*σ*∇*V*) = 0, where *σ* represents the electrical conductivity and *V* represents the electrical potential. Dirichlet boundary condition was set to fixed potentials imposed on one of the active electrodes (anterior electrode) and the return electrode. Then, the finite element solver GetDP (Dular et al., 1998) implemented in SimNIBS (Puonti et al., 2020) was used by employing the Galerkin method to calculate the electric field distribution inside the brain (Opitz et al., 2015). The injection current from each active electrode was assumed to be 1 mA. The same process was applied for the other active electrode (the posterior electrode) and the return electrode, resulting in two independent electric field distributions, **E**_**1**_ (anterior electrode – return electrode) and **E**_**2**_ (posterior electrode – return electrode).

Then, the direction of electric fields was captured in the anterior – posterior direction for each sEEG electrode. Considering the desired *d*-direction of electric fields for a specific sEEG electrode, resultant electric fields in this direction were defined as 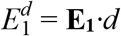 and 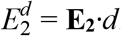. Then, we employed a phasor analysis to the resultant electric field distributions obtained from the previous step. Assuming that alternating currents at a specific frequency with phases of *θ*_1_ and *θ*_2_ were injected through the anterior and posterior electrodes, respectively, the phasic electric fields of *P*_1_ and *P*_2_ at any given location were determined as follows:

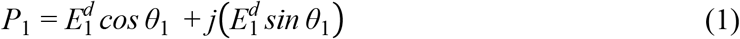

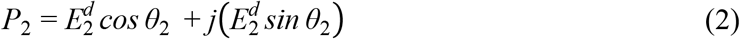

Then, the electric field *P* at any point generated from three-electrode tACS can be determined by the superposition of the two electric fields *P*_1_ and *P*_2_ as

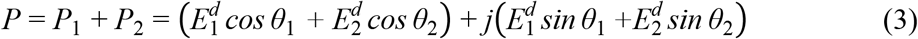

From the electric field *P*, we can estimate the amplitude 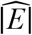 and the phase 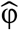 of the electric field *P* as follows:

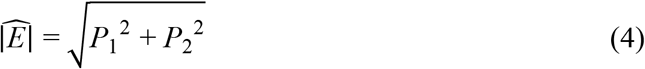

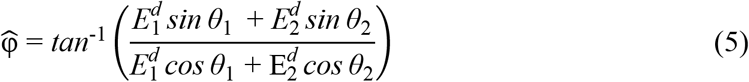

With this step, we can determine the phase and amplitude in the anterior-posterior direction for each contact of a specific sEEG electrode, followed by unwrapping of the phase angles and centering them between –π/2 and π/2. Given that *d*-direction was different among sEEG electrodes, we repeated the abovementioned steps for each sEEG electrode for each monkey to derive the accurate phase and amplitude at each electrode contact considering the directionality of electric fields as in *in vivo* experiments.

### 2.3. Data analysis

#### 2.3.1. Comparison between measured and simulated results

We visualized the distribution of the phase and amplitude of the directional electric fields obtained from the measurements and simulations for 22 stimulation conditions. Also, the polar graph with normalized amplitude for each stimulation condition was illustrated for each sEEG electrode. Then, we quantified the similarity between measured and simulated phase values by the circular correlation coefficient using the circular statistics toolbox (Jammalamadaka and SenGupta, 2001), as follows:

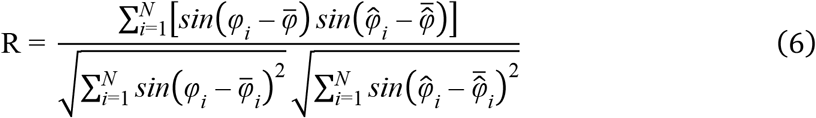

where *N* is the total number of contacts. φ_*i*_ and 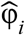 represent the phase of the *i*-th contacts in measurements and simulations, respectively. 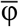 and 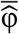 represent the mean values for these phases. For a comparison of the amplitude, we used Pearson’s correlation to determine the correlation coefficient between measured and simulated amplitudes. Since we already accounted for the directionality of each sEEG electrode, R values were calculated using all contacts.

#### 2.3.2. Effects of the return electrode placements

Once the direction of electric fields is determined to be anterior to posterior, the placement of the return electrode plays an important role in the characterization of electric fields. Thus, a small displacement of the return electrode in head models can cause a mismatch with the actual measurements. To explore this, the return electrode location was shifted in a 7-by-7 grid in the anterior-posterior (7 steps) and inferior-superior (7 steps) directions, with a step size of 5 mm (50% of the electrode radius), relative to the original location of the return electrode (which is referred to as the center of the grid), resulting in a total of 49 displacement points on the scalp. Certain points were excluded because they overlapped with the left ear, where the electrode could not be attached. For each displacement point, the simulated and measured results were compared.

#### 2.3.3. Optimization of electrical conductivity

In the current study, the electrical conductivity of human tissues was applied to the monkey models. This may cause a disparity between simulation and experiment, specifically for the amplitude of electric fields. To overcome this gap, we optimized the electrical conductivity by comparing measured and simulated amplitudes obtained from all stimulation conditions to find the best match. We employed the same optimization problem as in the previous study (Huang et al., 2017), as follows:

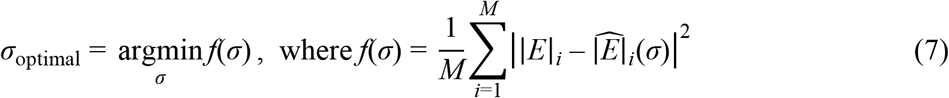

where |*E*|_*i*_ is the measured amplitude at the contact *i* and 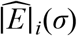 represents the simulated amplitude depending on the estimated electrical conductivity *σ*. *M* is the total number corresponding to the number of sEEG contacts × stimulation conditions. The problem was solved using a pattern search algorithm (Audet and Dennis Jr, 2002). This algorithm is suitable for finding the solution that has the lowest error value on discontinuous and nondifferential functions. The conductivity value of the original phasor analysis was used as the initial conductivity. *σ* was iteratively updated within a given range specified in the previous study (Huang et al., 2017) to minimize the cost function *f*(*σ*). The optimal electrical conductivity σ_optimal_ was determined for both monkeys. In the optimization process, we only considered the electrical conductivity of four tissues, including the scalp, skull, GM, and WM, as a variable, while the electrical conductivity of other liquid-filled tissues (CSF and eyes) is assumed to be a constant. We then investigated whether employing the optimal conductivity in simulations may effectively lessen the disagreement with *in vivo* recordings by calculating the absolute error for each stimulation condition as

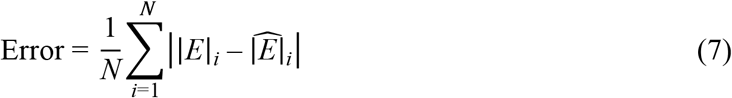

where *N* is the total number of sEEG electrode contacts. With either initial or optimal conductivity, we can derive the errors for two conductivity cases for the 22 different stimulation conditions, followed by a non-parametric statistical comparison of these two errors using the Wilcoxon signed-rank test.

## 3. Results

### 3.1. Comparison between simulated and measured results

**Figure 2** shows the comparison between the simulated and measured phase distributions and polar graph with normalized amplitude for sEEG electrode A2 for four representative stimulation conditions (45°, 135°, 225°, and 315°) in monkey 1. The phase gradient is well represented in the computational simulation relative to *in vivo* experiments (see **Fig. 2A**). Similarly, **Figure. 2B** shows simulated phase values are in good agreement with the measured phase values, with a relatively high electric field amplitude delivered to the posterior contacts in both simulations and experiments. The other polar graphs for the other sEEG electrodes for both monkeys are shown in **Supplementary Figures. 1-5**.

**Figure. 2.**
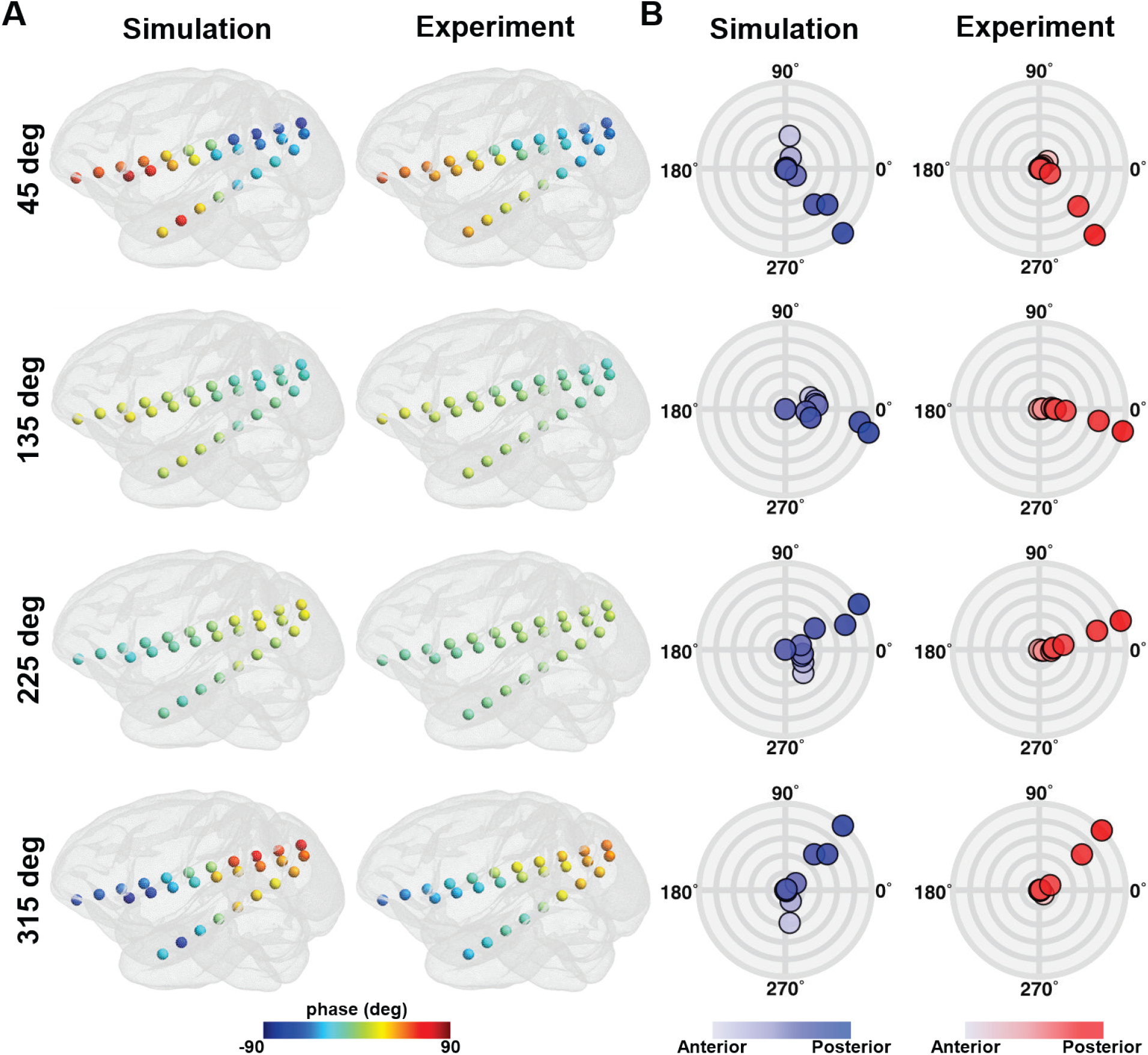
Comparison between simulations and *in vivo* measurements for four representative stimulation conditions (45°, 135°, 225°, and 315°). A) The phase distribution at all sEEG electrodes in monkey 1. B) The polar graph with the phase and the normalized amplitude at all contacts of sEEG electrode A2 in monkey 1. Colored circles represent individual contacts in the sEEG electrode, with a gradual color gradient along the anterior-posterior direction. The outermost circular line in the polar graph represents the normalized amplitude of 1, with an interval of 0.2 between circular lines.

We confirmed that the simulation can predict the phase distribution for both monkeys (**Fig. 3A and B**) across all stimulation conditions. Note that the phase and amplitude distributions at sEEG electrodes for the other stimulation conditions are illustrated in **Supplementary Figures. 6 and 7**. To quantify the similarity, we calculated the correlation coefficient (R) for all stimulation conditions. For instance, we illustrate a correlation graph for the 45° stimulation condition, showing simulated phases are in good agreement with measured phases (**Fig. 3C and D, left panel**). Likewise, under most stimulation conditions, the R value is close to 0.9, with the mean R values of 0.89 and 0.90 for monkey 1 and monkey 2, respectively (**Fig. 3C and D, right panel**). **Figure 4** shows the comparison between the simulated and measured amplitude distributions. Amplitude distributions in all sEEG electrodes are quite similar in both the simulation and experiment; however, the amplitude is comparably greater in the simulations for both monkeys (**Fig. 4A and B**). This is also evident in the correlation graph with higher values on the axis representing simulations (**Fig. 4C and D**). Nevertheless, the simulation accurately predicts the distribution of the amplitudes for the *in vivo* experiments with the mean R values of 0.81 and 0.75 for monkey 1 and monkey 2, respectively.

**Figure. 3.**
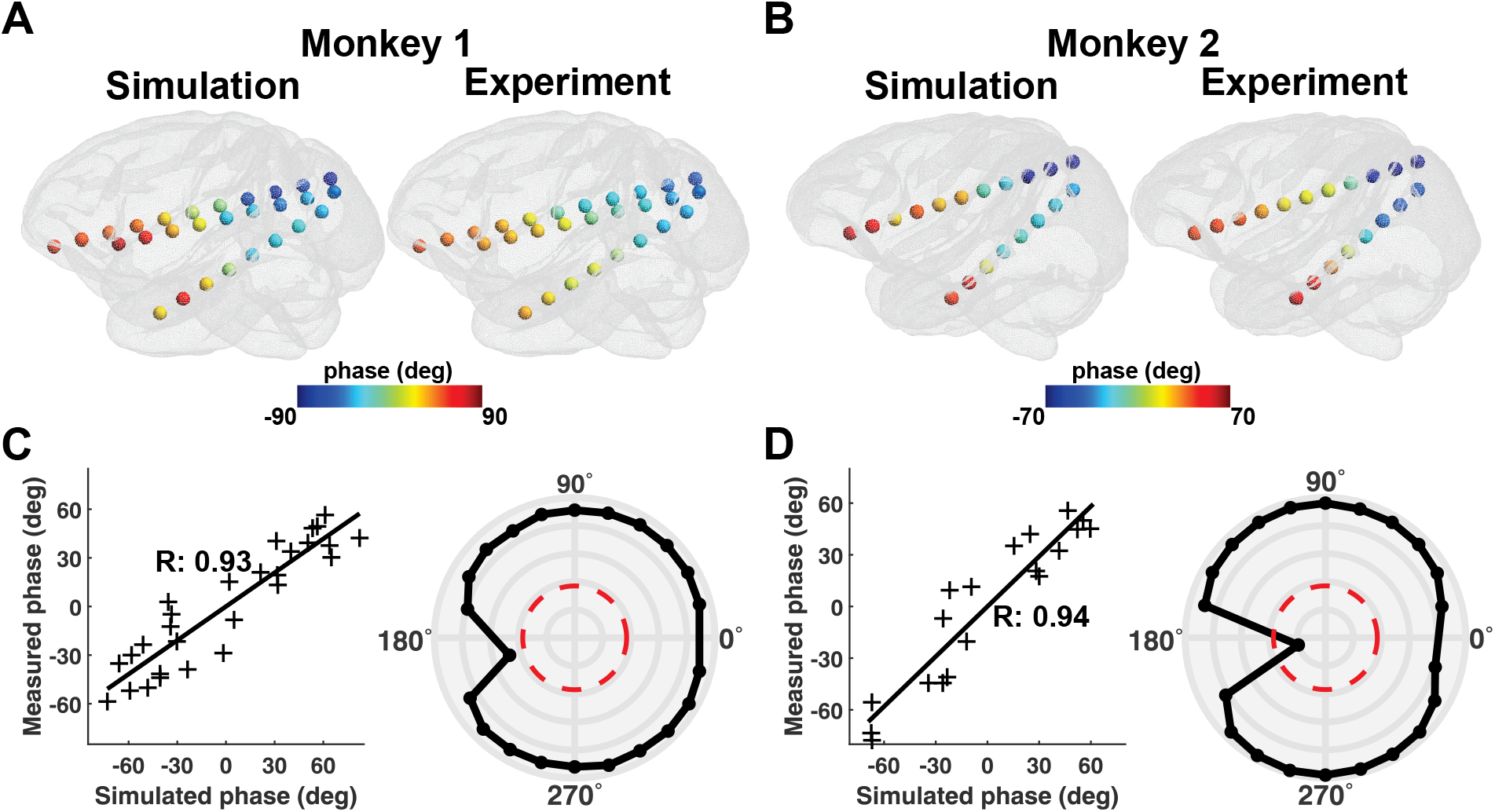
Comparison between simulations and *in vivo* measurements for the phase. A, B) Illustration of the phase distribution at all sEEG electrodes during 45° stimulation condition for both monkeys. C, D) The example correlation between simulated and measured phases for 45° stimulation condition (left panel). The polar graph depicts the correlation values between measurements and simulations for each stimulation condition for both monkeys (right panel). The red line in the polar graph denotes the significance level of *p* = 0.05, while the outermost circular line in the polar graph represents a correlation value of 1, with an interval of 0.2 between circular lines.

**Figure. 4.**
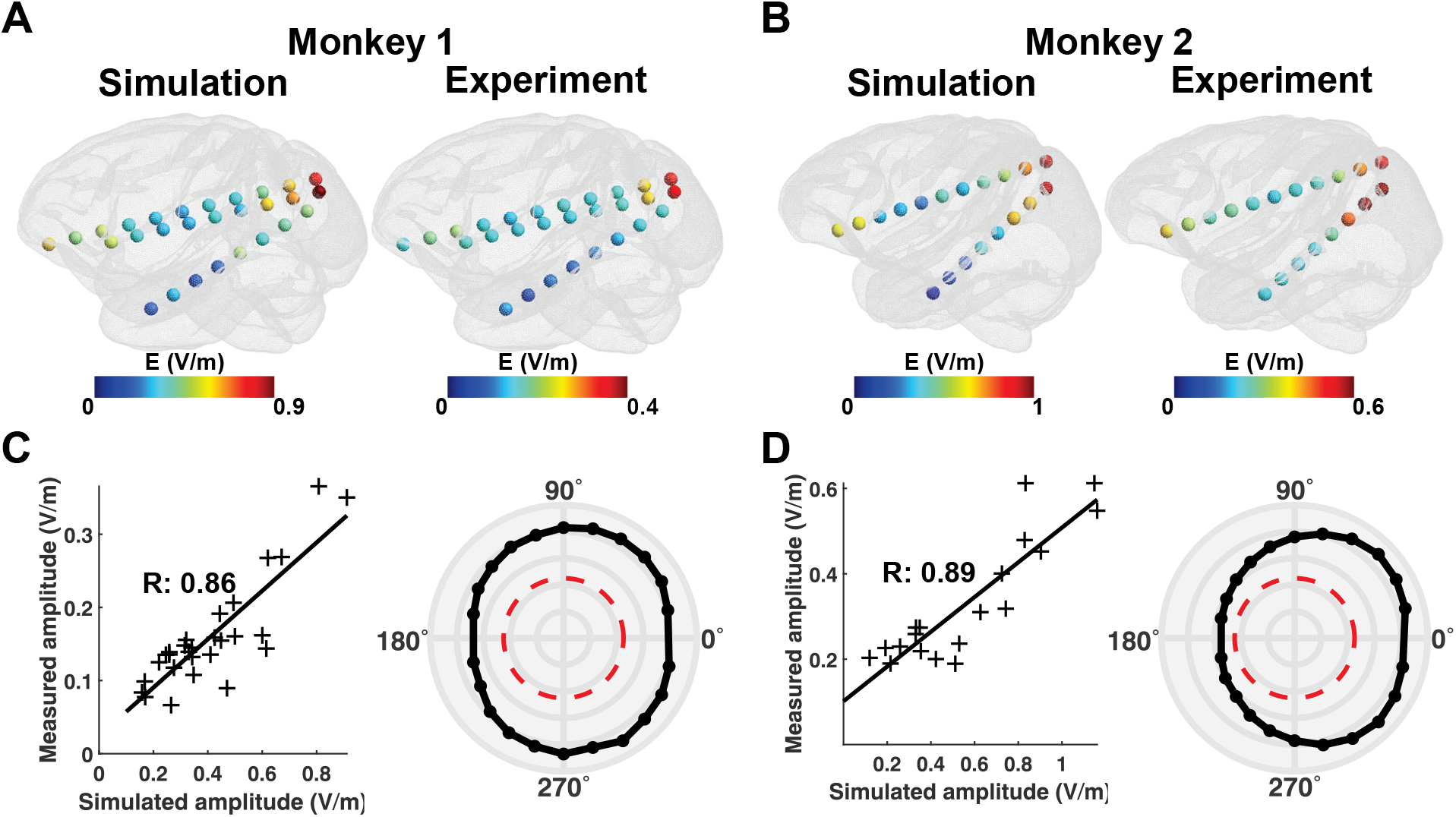
Comparison between simulations and *in vivo* measurements for the amplitude. A, B) Illustration of the amplitude distribution at all sEEG electrodes during the 45° stimulation condition for both monkeys. C, D) The example correlation between simulated and measured amplitudes for 45° stimulation condition (left panel). The polar graph depicts the correlation values for each stimulation condition for both monkeys (right panel). The red line in the polar graph denotes the significance level of *p* = 0.05, while the outermost circular line in the polar graph represents a correlation value of 1, with an interval of 0.2 between circular lines.

### 3.1. Change in the correlation depending on the return electrode placement

Next, we examined how a small displacement of the return electrode in head models affects the correlation between simulated and measured outcomes. For both monkeys, the mean correlation coefficient of the phase and amplitude for all stimulation conditions was highest close to the original location (**Figs. 5A and C**). However, the correlations continuously declined as the return electrode was moved farther from the center. For monkey 1, the correlation values did not markedly change when moving the return electrode in the inferior-superior direction, although a considerable decrease in both correlation values of the phase and amplitude was observed in the anterior-posterior direction as shown in **Fig. 5B**. In monkey 2, the correlation values decreased regardless of the displacement direction as the return electrode was attached far from the center (**Fig. 5D**). Furthermore, in both monkeys, the correlation values of amplitudes are considerably more sensitive to the return electrode shifts than the phase correlations. The correlation values tend to be higher within the range between -5 mm and 5 mm from the center, implying that the minimum radius of 5 mm (50% of the electrode radius) for the return electrode displacement is most optimal for an accurate simulation to capture phasic electric field distribution during a tACS experiment.

**Figure. 5.**
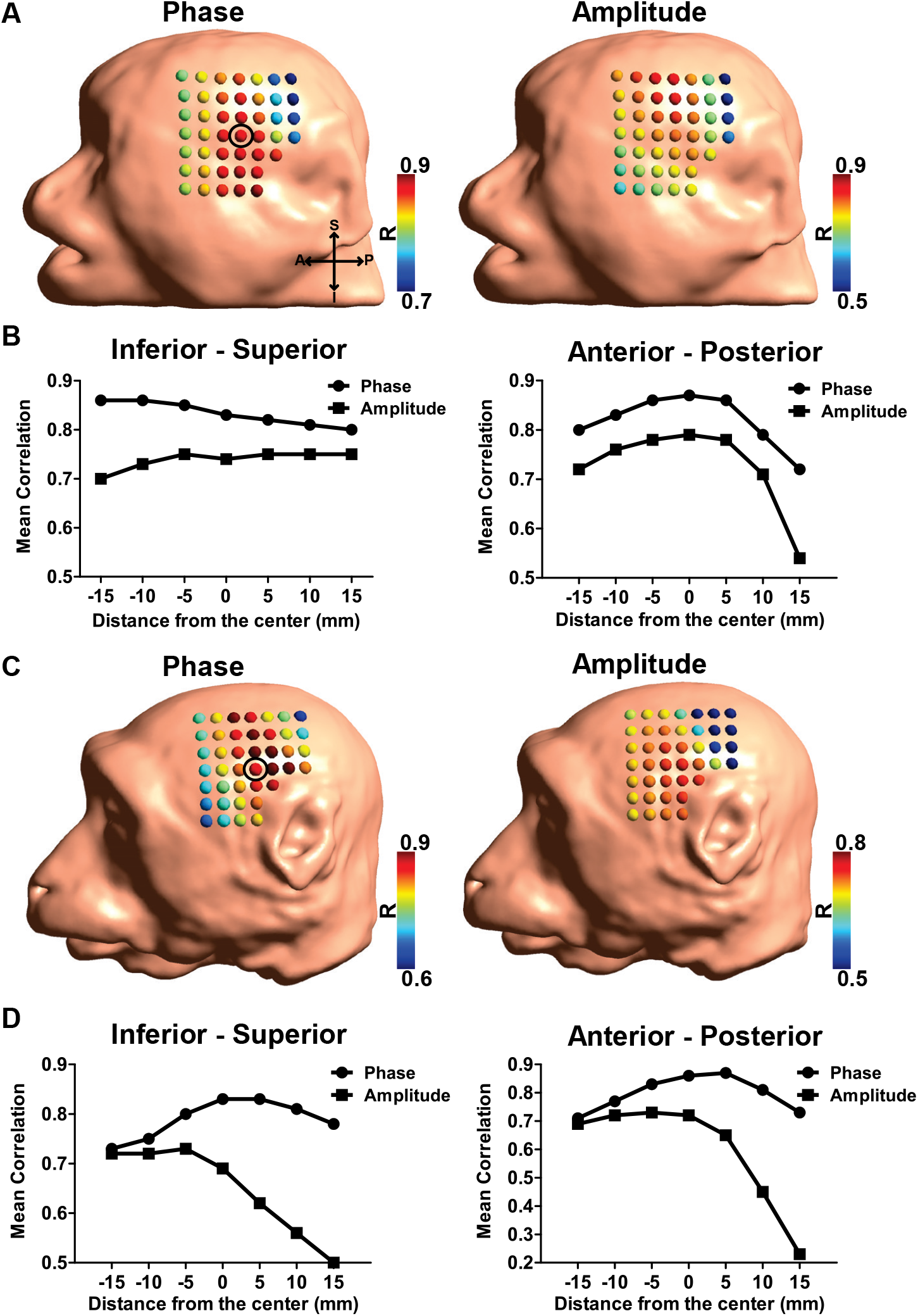
Effects of a small displacement of the return electrode. The mean correlation values between simulated and measured results (phase and amplitude) for all stimulation conditions as the return electrode was moved in steps of 5 mm for A) monkey 1 and C) monkey 2. The black circle indicates the center of the displacement (the original position of the return electrode), while the dots in head models represent the return electrode locations. B, D) Mean correlation values for the phase and amplitude with a distance from the center along the inferior-superior (left panel) and anterior-posterior directions (right panel).

### 3.1. Effects of employing optimal electrical conductivity

Given a comparable difference in the amplitude between simulations and experiments, we individually calibrated the electrical conductivities in head models to optimize the similarity in amplitudes. The correlation values are either consistent with or slightly higher than the standard conductivity values used in computational modeling previously (Alekseichuk et al., 2019b) for the 22 stimulation conditions when employing the optimal conductivity, whereas the amplitude itself was decreased for both monkeys in comparison to those employing the initial conductivity (**Figs. 6A and B**). We confirmed that the mean absolute errors in the case of the optimal conductivity (0.05 ± 0.01 for monkey 1 and 0.10 ± 0.02 for monkey 2) were significantly smaller than those in the case of the initial conductivity of 0.26 ± 0.01 and 0.27 ± 0.02, respectively (*p* < 0.05 for both monkeys). This indicates that an optimization process can be used to determine the tissue’s electrical properties, thereby overcoming the difference of the amplitude between the computational simulation and *in vivo* experiments. The optimal conductivity values are listed in **Supplementary Table. 1**.

**Figure. 6.**
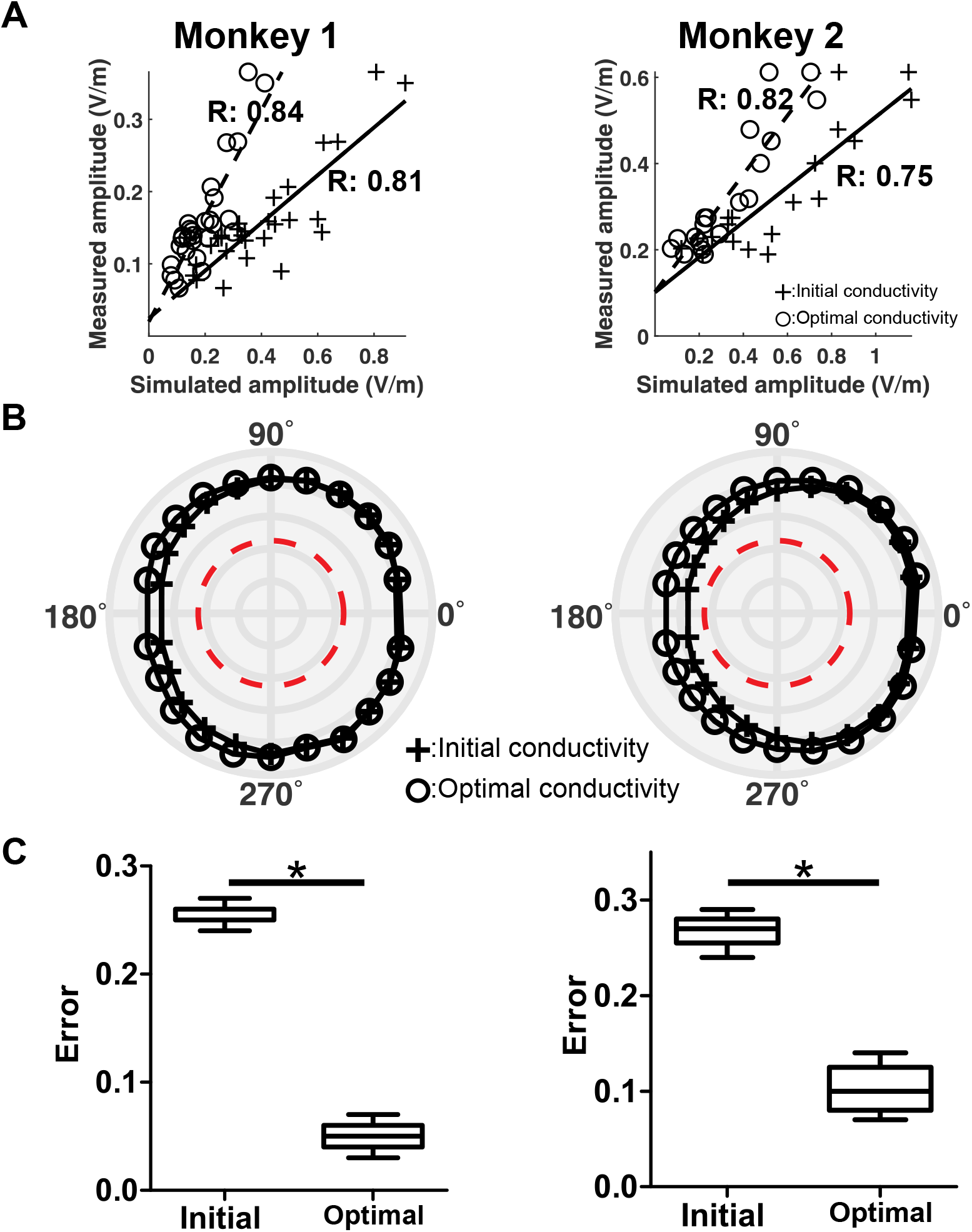
Effects of employing the optimal conductivity in the computational simulations for monkey 1 (left column) and monkey 2 (right column). A) Comparison of the correlation of the amplitude with that obtained by applying the optimal conductivity. The cross symbols with the solid line are associated with the simulated amplitude obtained by applying the initial conductivity, and the circle symbols with the dotted line are associated with the simulated amplitude when applying the optimal conductivity. B) Correlation values for all 22 stimulation conditions when either applying initial or optimal conductivities. The red line represents the significance level of *p* = 0.05. C) The absolute error for all stimulation conditions (*n* = 23) between measured and simulated amplitudes obtained by either applying the initial or optimal conductivities (\**p* < 0.05).

## 4. Discussion

This is the first study to validate the accuracy of phasor simulation for tACS with *in vivo* measurements. Our study has three main findings: i) the phasor simulation using head models achieves an accurate prediction of the phase distribution inside the brain for multi-channel tACS; ii) the simulation precisely predicts the phase in *in vivo* experiments only when the return electrode is positioned within a small radius from the actual location; and iii) the tendency of the simulated electric field amplitude distribution follows the measured amplitude, while an overestimation in the electric field amplitude can be calibrated by optimizing the electrical conductivity. Notably, 22 experimental stimulation conditions for tACS in two monkeys were used for our analysis. We further developed individual head models that can adequately capture the anatomical structure. Consequently, our findings provide solid and clear evidence that phasor analysis in the head model can capture the properties of tACS electric fields from phase-shifted inputs.

Our findings show that the phasor simulation accurately predicts the phase gradient along sEEG electrodes as observed in intracranial recordings (**Supplementary Figures 6A and 7A**). We confirmed that a phase shift between the anterior and posterior electrodes is visible in simulations when injecting the currents with a phase difference higher than 180° (Alekseichuk et al., 2019a). As depicted in **Fig. 3**, under most stimulation conditions, the correlation values between simulated and measured phases were between 0.9 and 0.95, with a maximum of 0.94 and 0.98, for monkey 1 and monkey 2, respectively (**Supplementary Figures 8A and 10A**). Still, some caution should be made when interpreting our findings. First, some differences exist in the phase value at each sEEG contact between simulations and measurements. This might be due to several factors, such as limited data quality of *in vivo* recordings at some contacts and the misestimation of electric fields due to uncertainty in the electrical properties of brain tissues. However, for traveling waves the phase difference (or phase gradient) across targeted regions is more important than the accurate phase estimation at a single location in the brain. From this perspective, our phase estimation can accurately capture the phase gradient in *in vivo* measurements. Thus, our findings provide important evidence that computational models based on phasor algebra can accurately estimate the phase of recorded electric fields extracted from the complex fourier values in FFT.

The location of the stimulation electrodes, especially the return electrode, is crucial for an accurate estimation of traveling wave electric fields. This is because the electric field direction will be determined as in-phasic or anti-phasic based on the location of the return electrode when the directionality of the electric field is determined (**Figure 1A**). Our results show that the return electrode has to be placed very precisely (< 5 mm distance) to properly estimate the phasic information of electric fields as in *in vivo* recordings. This is in line with a previous study suggesting that the minimal displacement required to ensure the accuracy of simulations in human head models was less than 10 mm (Opitz et al., 2018). Furthermore, the location of the return electrode is related to the direction of the electric field. **Figure 5** shows that correlation values are more sensitive to the anterior-posterior displacement of the return electrode. Especially, the correlation values for the amplitude dropped dramatically when the return electrode was attached to the posterior part of the head. This could be explained by a previous finding indicating that electric fields are predominantly shunted through the scalp when the electrodes are close together (Faria et al., 2011; Seibt et al., 2015). Therefore, the precise location of the return electrode while considering the direction of the traveling wave to be manipulated by tACS is essential for a reliable prediction.

The phasor simulation also estimated the spatial distribution of electric field amplitude inside the brain at a similar level to previous validation studies (Huang et al., 2017; Opitz et al., 2016; Puonti et al., 2020) (**Supplementary Figures. 6B and 7B**). However, the simulated amplitude itself was considerably higher than the measured one. This is due to a systematical overestimation in head models when using a well-known electrical property measured in ex-vivo conditions (Huang et al., 2017; Opitz et al., 2017; Opitz et al., 2018). For instance, the maximum amplitudes in simulations were about 0.8 V/m and 1.2 V/m for monkey 1 and monkey 2, respectively, whereas those in measurements were about 0.4 V/m and 0.8 V/m (**Supplementary Figures. 9A and 11A**). This difference can be significantly reduced by using optimal conductivity values in simulations. Despite this, we must be cautious in interpreting optimization results. The aim of the optimization was to minimize the error between simulated and measured amplitudes. Thus, it is more appropriate to regard the role of optimal conductivity as a calibration (Huang et al., 2017).

NHP models have been commonly used to explore biophysical effects of tACS due to their similarity of anatomical structures to humans (Huang et al., 2021; Johnson et al., 2020; Opitz et al., 2016). Therefore, we can extend our understanding of our results to human participants. It will be important to place the return electrode precisely in human experiments. This can be achieved, for example, by digitizing the coordinates of electrodes on the scalp corresponding to those in the human head model using a 3D digitizer in clinical experiments (Nieminen et al., 2022). In addition, the phase values at sEEG contacts are not markedly different between the cases employing initial and optimal conductivities. This indicates that the phase distribution is quite robust to the change of tissue conductivities (**Supplementary Figures 8 - 11**). Our results indicate that ohmic properties dominantly affect the amplitude of oscillations, not phasic information. Thus, a consistent phase distribution would be expected among participants as long as the stimulation phase condition is the same. As ongoing brain oscillations have their own time-lag patterns, it will be important to generate a phase gradient that is optimally aligned to this pattern. To do this, it is required to develop a multi-channel tACS system with optimization that can determine optimal electrode conditions (e.g., the phase and amplitude of injecting currents, electrode position) to generate the desired phase gradient over targeted regions.

In summary, we validated the accuracy of the phasor simulation in two monkeys via *in vivo* measurements. Our findings provide clear evidence that the phasor simulation can accurately estimate the phase distribution in the form of traveling waves as well as the spatial distribution of the amplitude inside the brain during multi-channel tACS. An additional calibration through the optimization of the electrical conductivity was required to better match predicted and measured electric field amplitudes. Our study lays the foundation for optimized multi-channel tACS that can manipulate ongoing brain oscillations in a phase-specific manner.

## Data Availability

Please contact the corresponding author for data requests.

## Author Contributions

SL and AO conceived the study and designed the experiments. SS advised on data analysis. IA, GL, CES, and AYF acquired and preprocessed the data. IA and NP conducted the head modeling. AO supervised the study.

## Acknowledgments

This research was supported in part by NIH (1RF1MH124909-01) and in part by a grant of the Korea Health Technology R&D Project through the Korea Health Industry Development Institute (KHIDI), funded by the Ministry of Health & Welfare, Republic of Korea (grant number: HI21C1234).

