## Supplementary materials for "Experimental validation of computational models for the prediction of phase distribution during multi-channel transcranial alternating current stimulation"

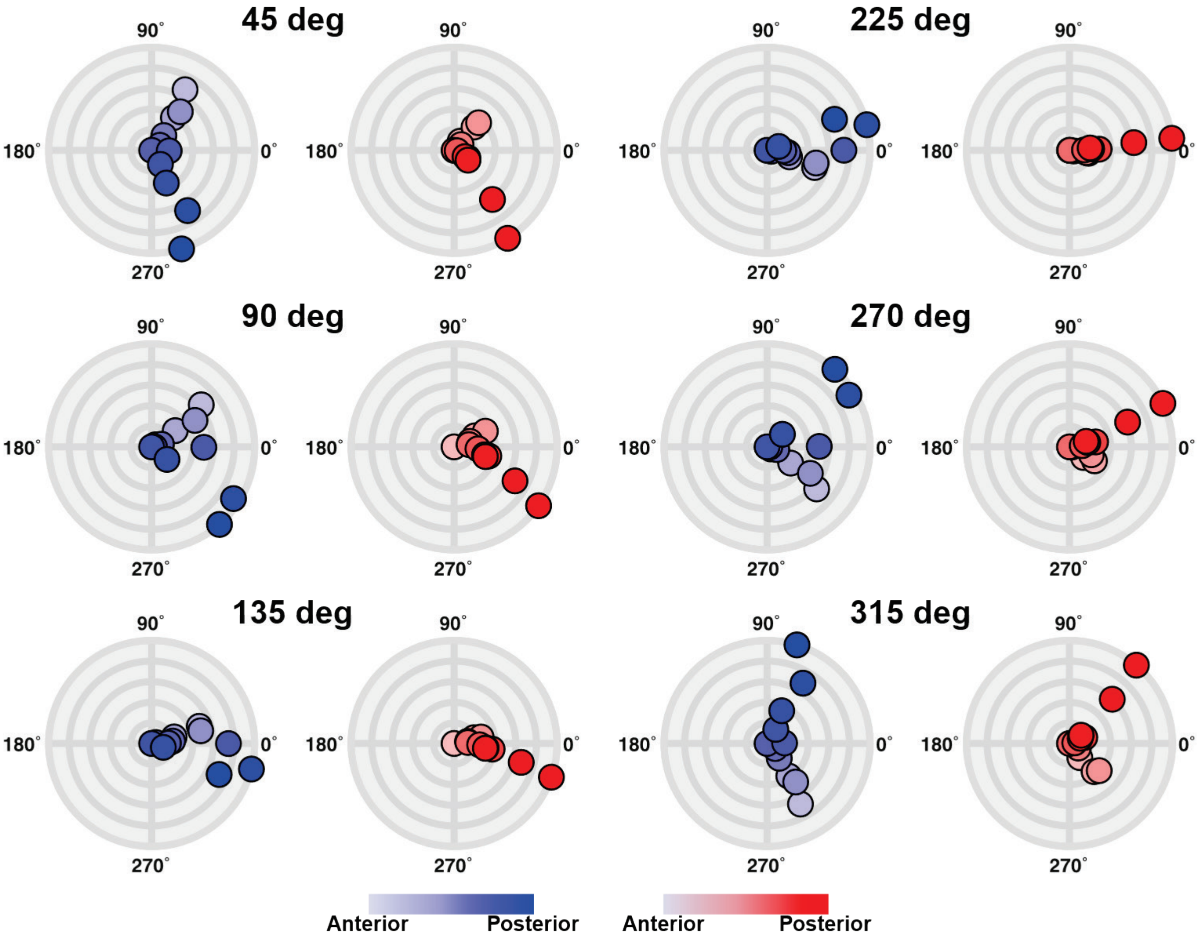


**Supplementary Figure. 1.** Illustration of polar graphs with the normalized amplitude for simulations (left, blue circles) and experiments (right, red circles) at sEEG electrode A1 in monkey 1. Six stimulation conditions are representatively depicted in the figure. Colored circles represent individual contacts in the sEEG electrode, with a gradual color gradient along the anterior-posterior direction. The outermost circular line in the polar graph represents the normalized amplitude of 1, with an interval of 0.2 between circular lines.

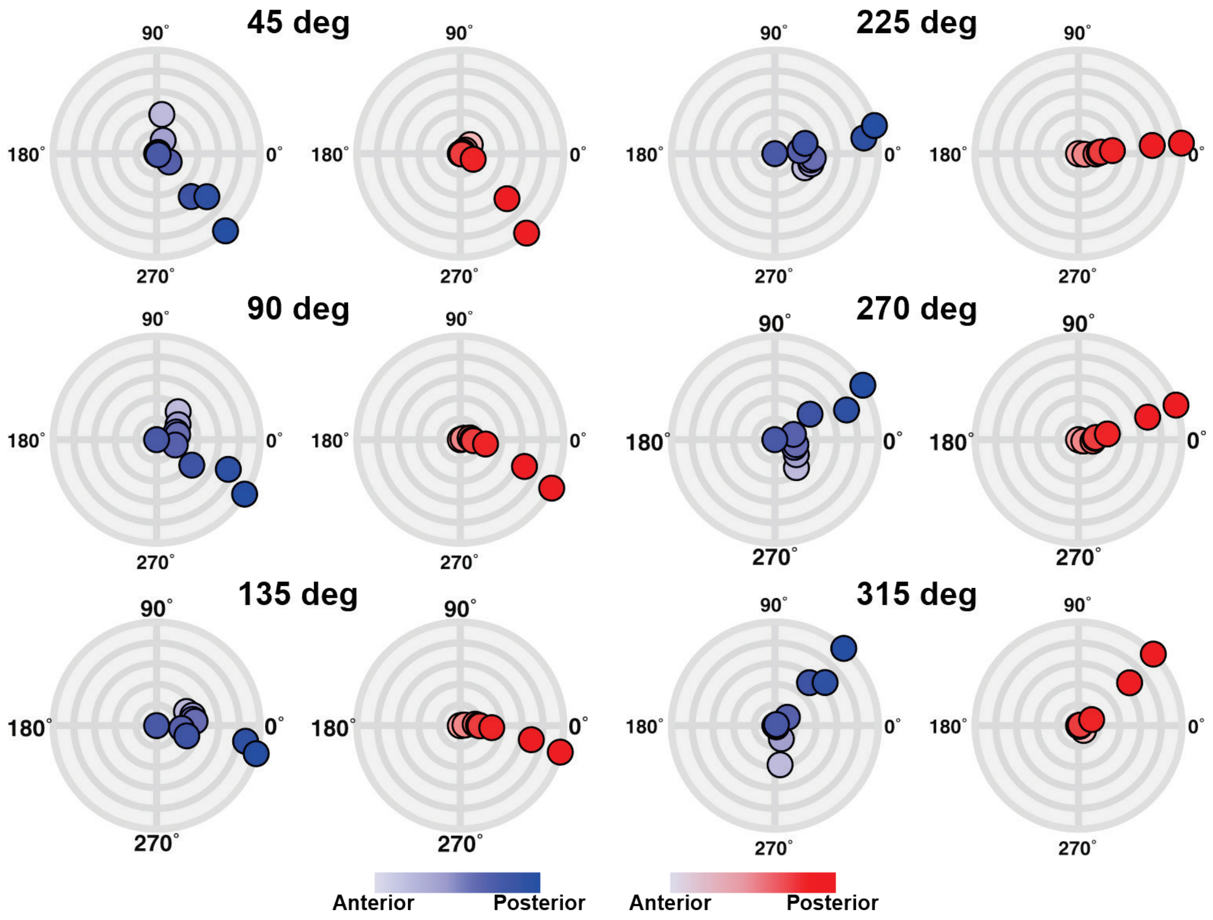


**Supplementary Figure. 2.** Illustration of polar graphs with the normalized amplitude for simulations (left, blue circles) and experiments (right, red circles) at sEEG electrode A2 in monkey 1. Six stimulation conditions are representatively depicted in the figure. Colored circles represent individual contacts in the sEEG electrode, with a gradual color gradient along the anterior-posterior direction. The outermost circular line in the polar graph represents the normalized amplitude of 1, with an interval of 0.2 between circular lines.

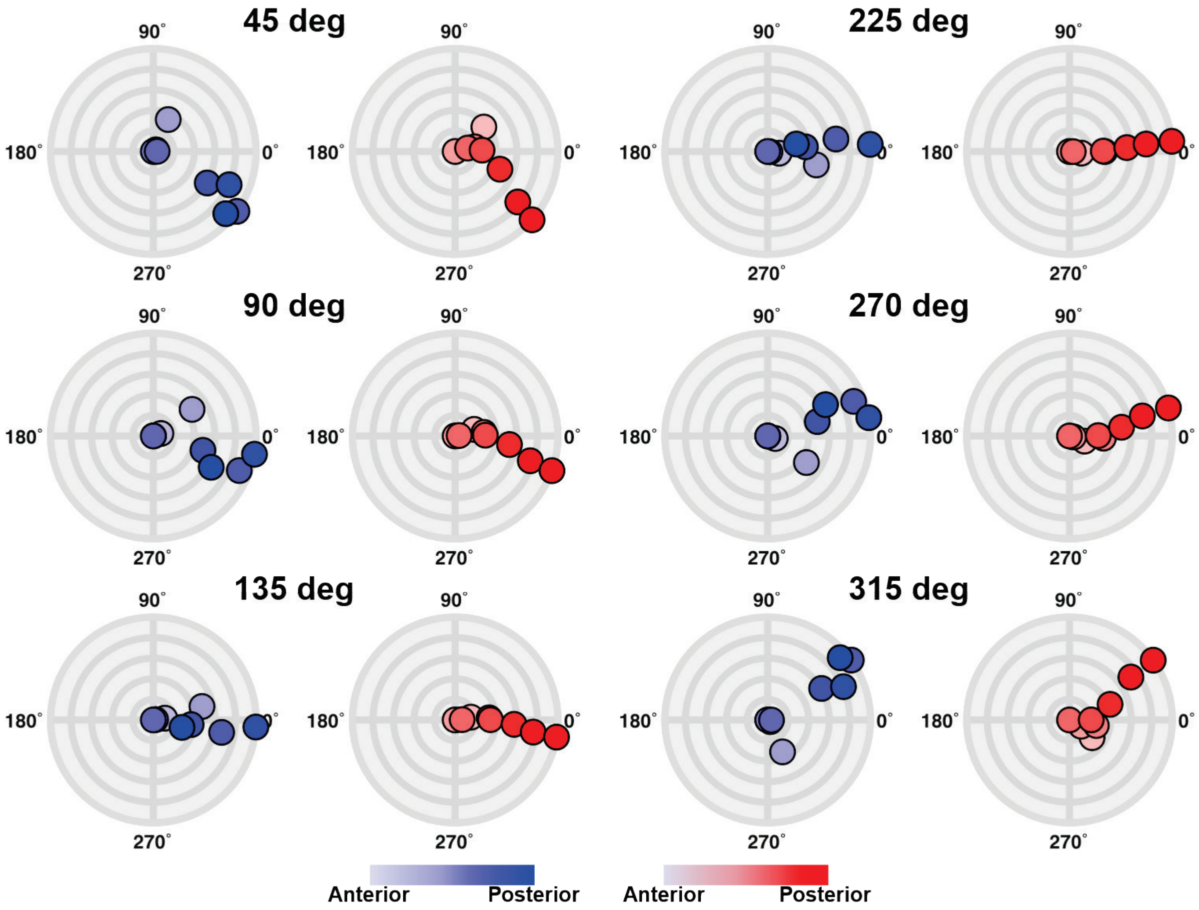


**Supplementary Figure. 3.** Illustration of polar graphs with the normalized amplitude for simulations (left, blue circles) and experiments (right, red circles) at sEEG electrode A3 in monkey 1. Six stimulation conditions are representatively depicted in the figure. Colored circles represent individual contacts in the sEEG electrode, with a gradual color gradient along the anterior-posterior direction. The outermost circular line in the polar graph represents the normalized amplitude of 1, with an interval of 0.2 between circular lines.

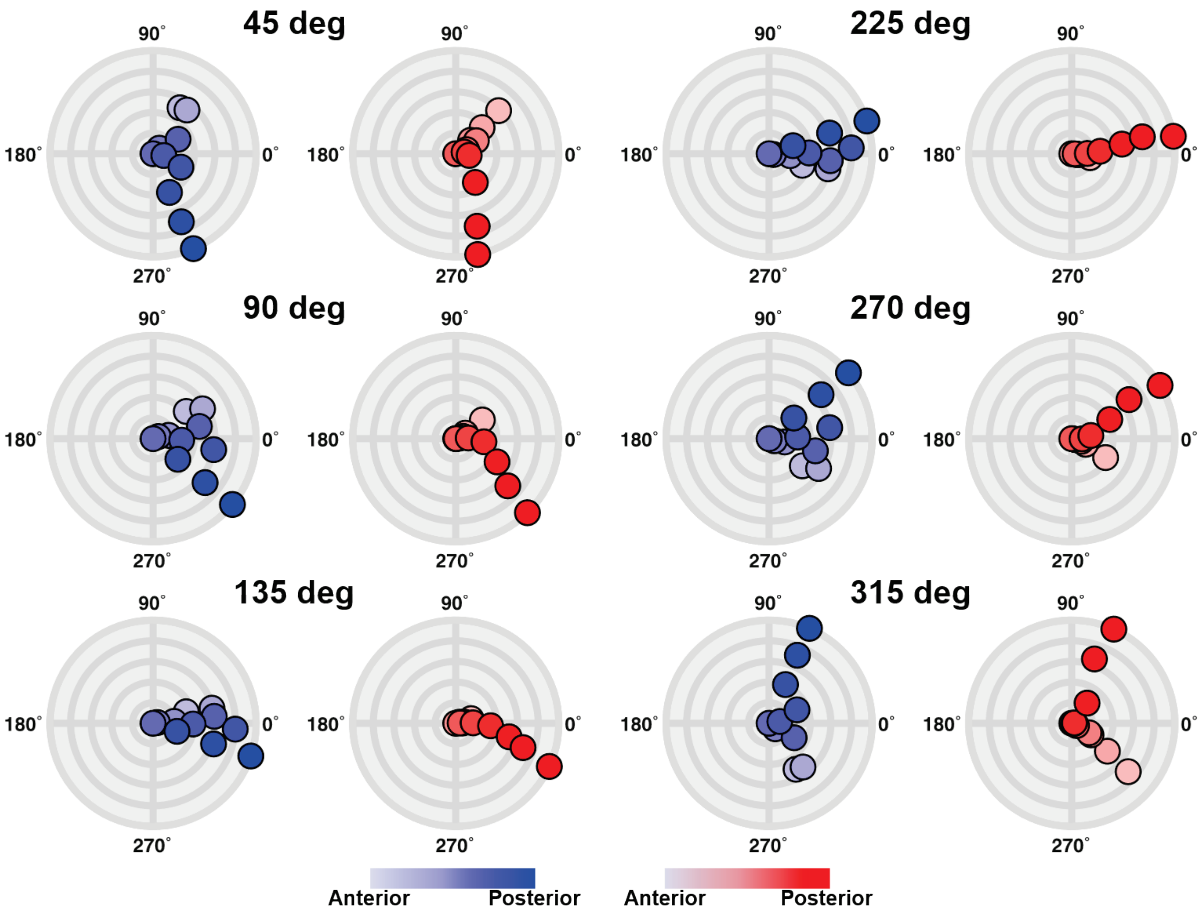


**Supplementary Figure. 4.** Illustration of polar graphs with the normalized amplitude for simulations (left, blue circles) and experiments (right, red circles) at sEEG electrode B1 in monkey 2. Six stimulation conditions are representatively depicted in the figure. Colored circles represent individual contacts in the sEEG electrode, with a gradual color gradient along the anterior-posterior direction. The outermost circular line in the polar graph represents the normalized amplitude of 1, with an interval of 0.2 between circular lines.

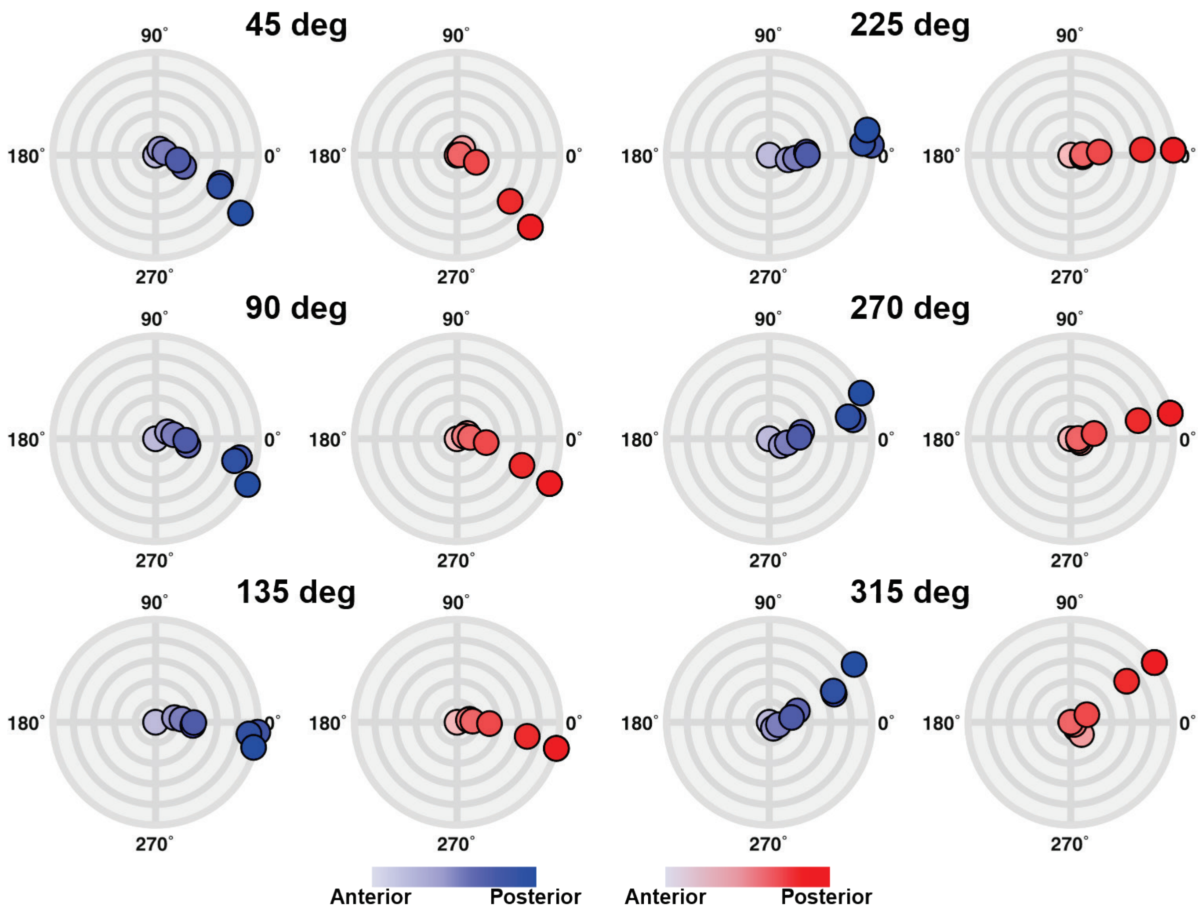


**Supplementary Figure. 5.** Illustration of polar graphs with the normalized amplitude for simulations (left, blue circles) and experiments (right, red circles) at sEEG electrode B2 in monkey 2. Six stimulation conditions are representatively depicted in the figure. Colored circles represent individual contacts in the sEEG electrode, with a gradual color gradient along the anterior-posterior direction. The outermost circular line in the polar graph represents the normalized amplitude of 1, with an interval of 0.2 between circular lines.

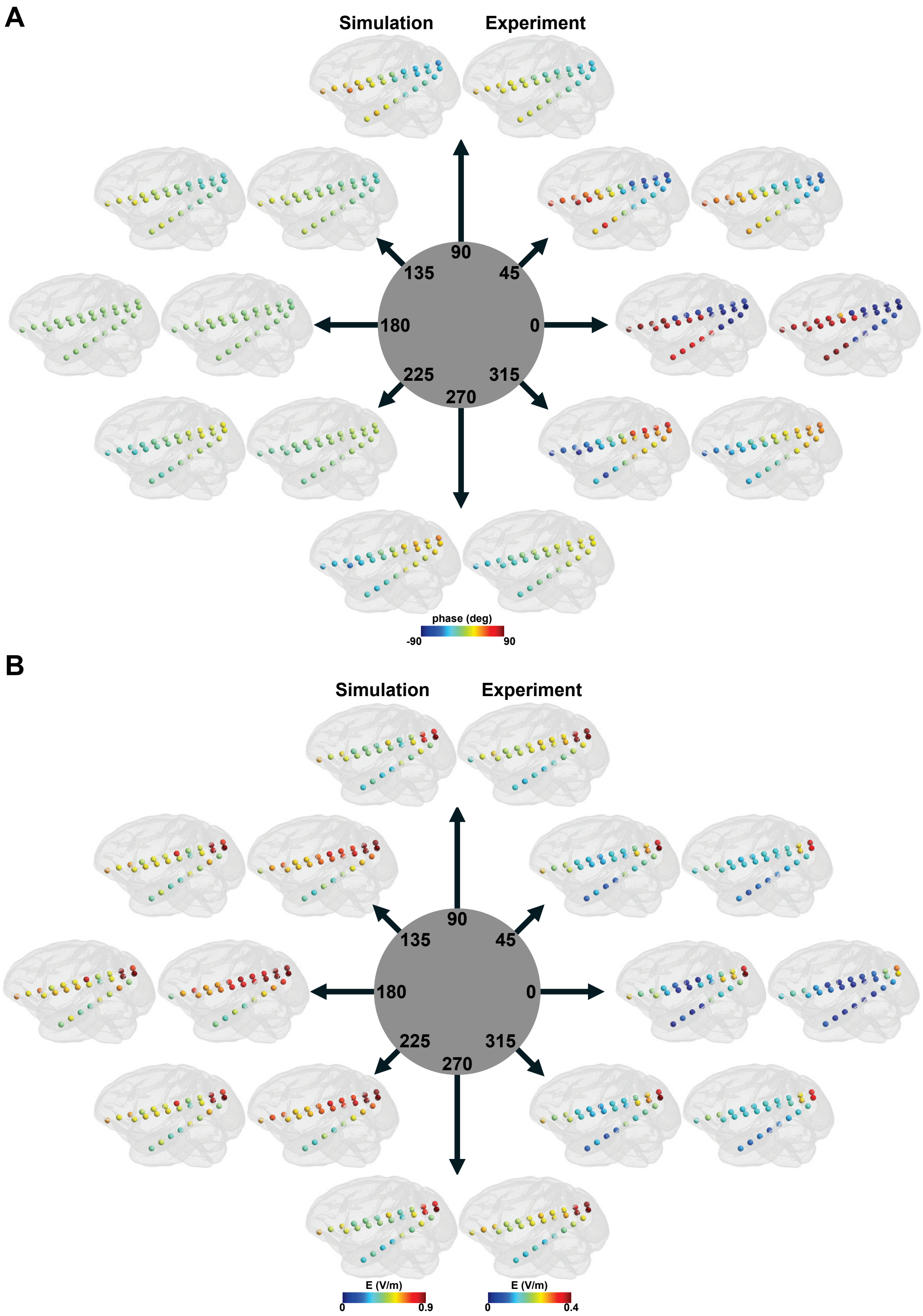


**Supplementary Figure. 6.** Comparison between simulations and *in vivo* experiments for A) the phase distribution and B) the amplitude distribution during multi-channel tACS for monkey 1. The number within the circle in the center represents the stimulation condition.

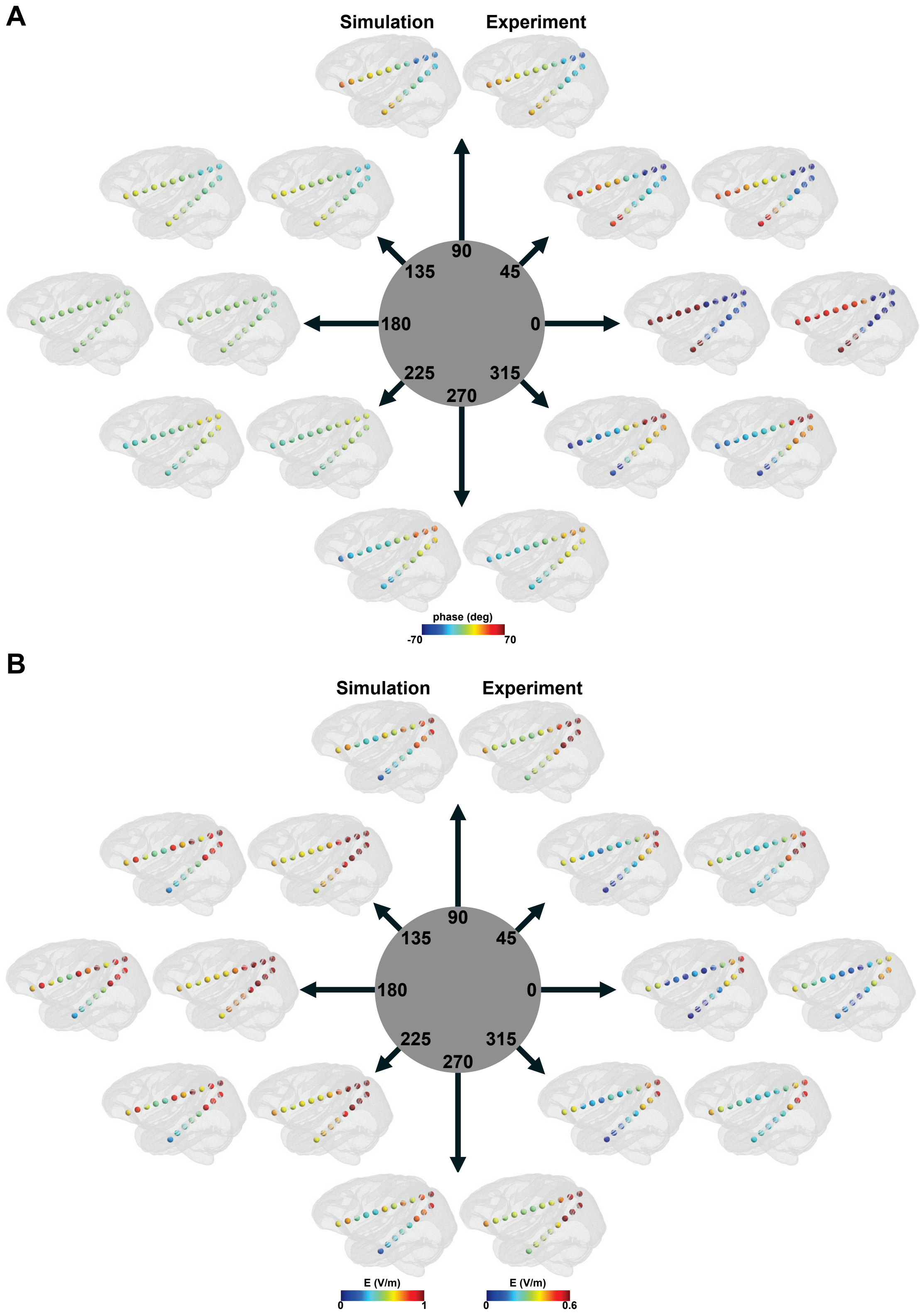


**Supplementary Figure. 7.** Comparison between simulations and *in vivo* experiments for A) the phase distribution and B) the amplitude distribution during multi-channel tACS for monkey 2. The number within the circle in the center represents the stimulation condition.

**Supplementary Table 1.** Optimal conductivity values for monkey 1 and monkey 2.

| | Monkey 1 | Monkey 2 |
|---|---|---|
| Scalp | 0.99 | 0.54 |
| Skull | 0.01 | 0.01 |
| GM | 0.37 | 0.58 |
| WM | 0.22 | 0.39 |

Unit: S/m

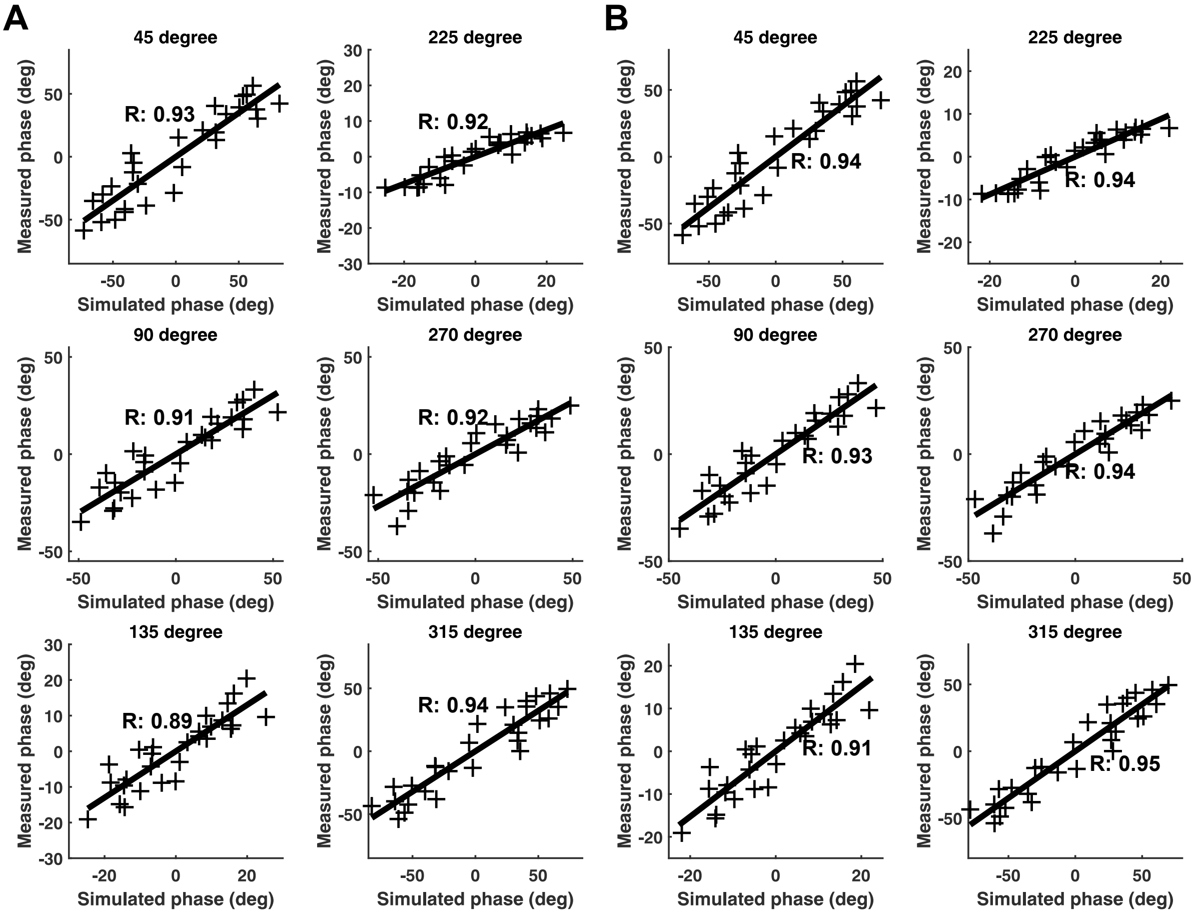


**Supplementary Figure. 8.** Correlation between the measured phase and either A) the simulated phase obtained when applying the initial conductivity or B) the simulated phase calculated with the optimal conductivity for monkey 1.

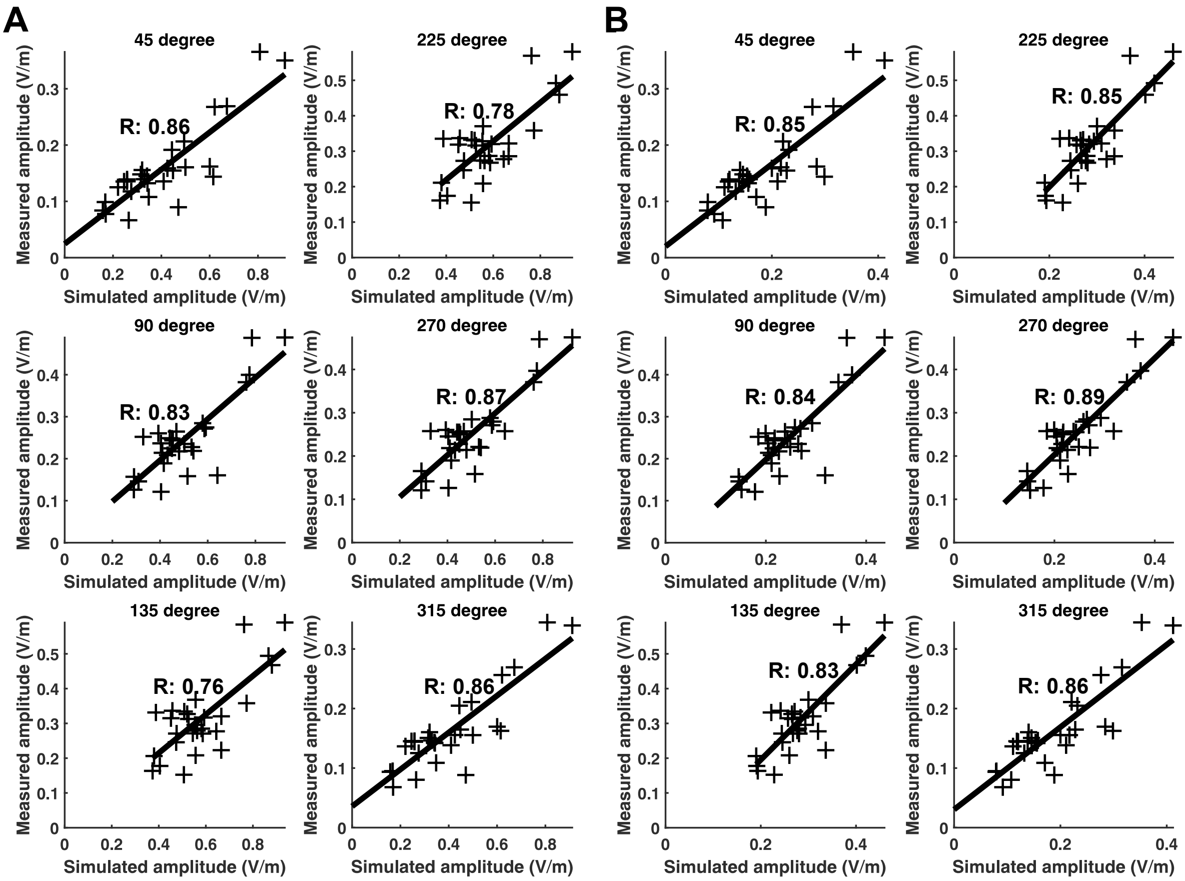


**Supplementary Figure. 9.** Correlation between the measured amplitude and either A) the simulated amplitude obtained when applying the initial conductivity or B) the simulated amplitude calculated with the optimal conductivity for monkey 1.

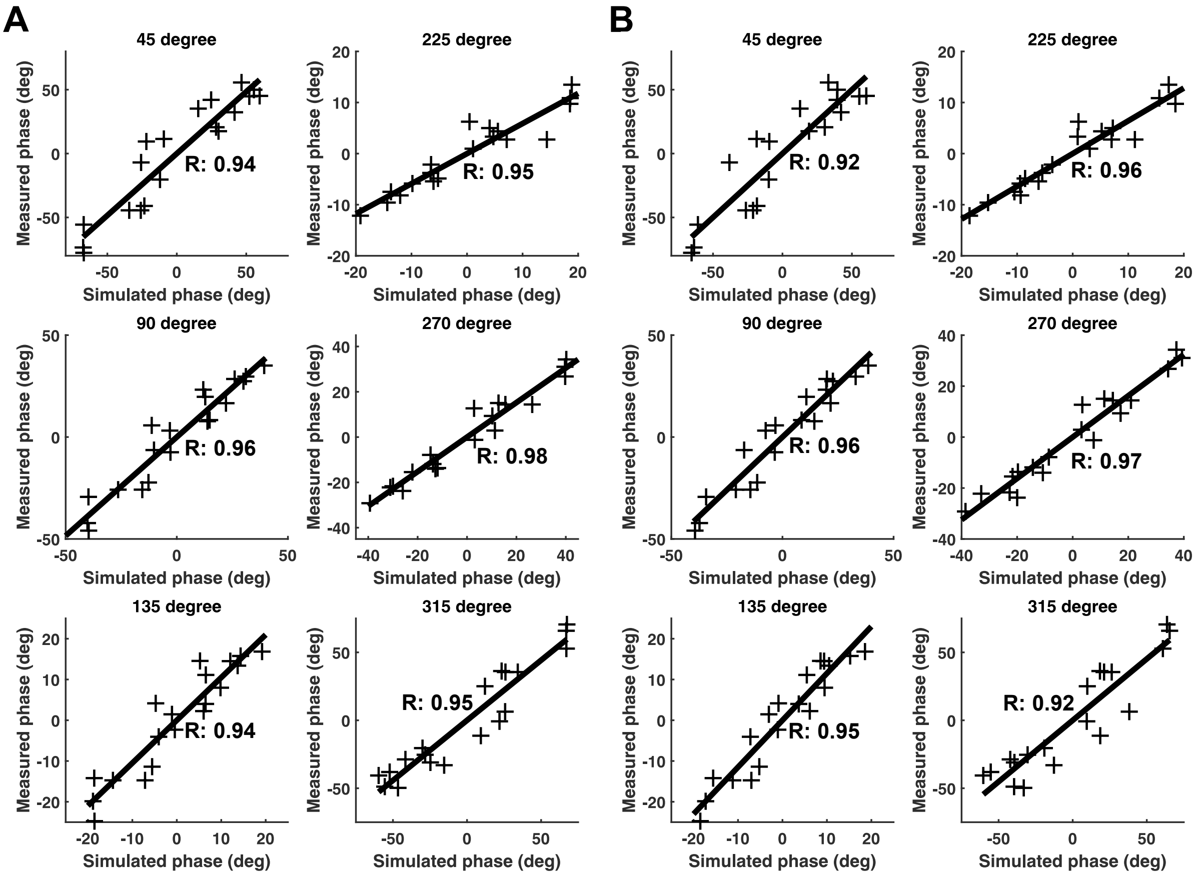


**Supplementary Figure. 10.** Correlation between the measured phase and either A) the simulated phase obtained when applying the initial conductivity or B) the simulated phase calculated with the optimal conductivity for monkey 2.

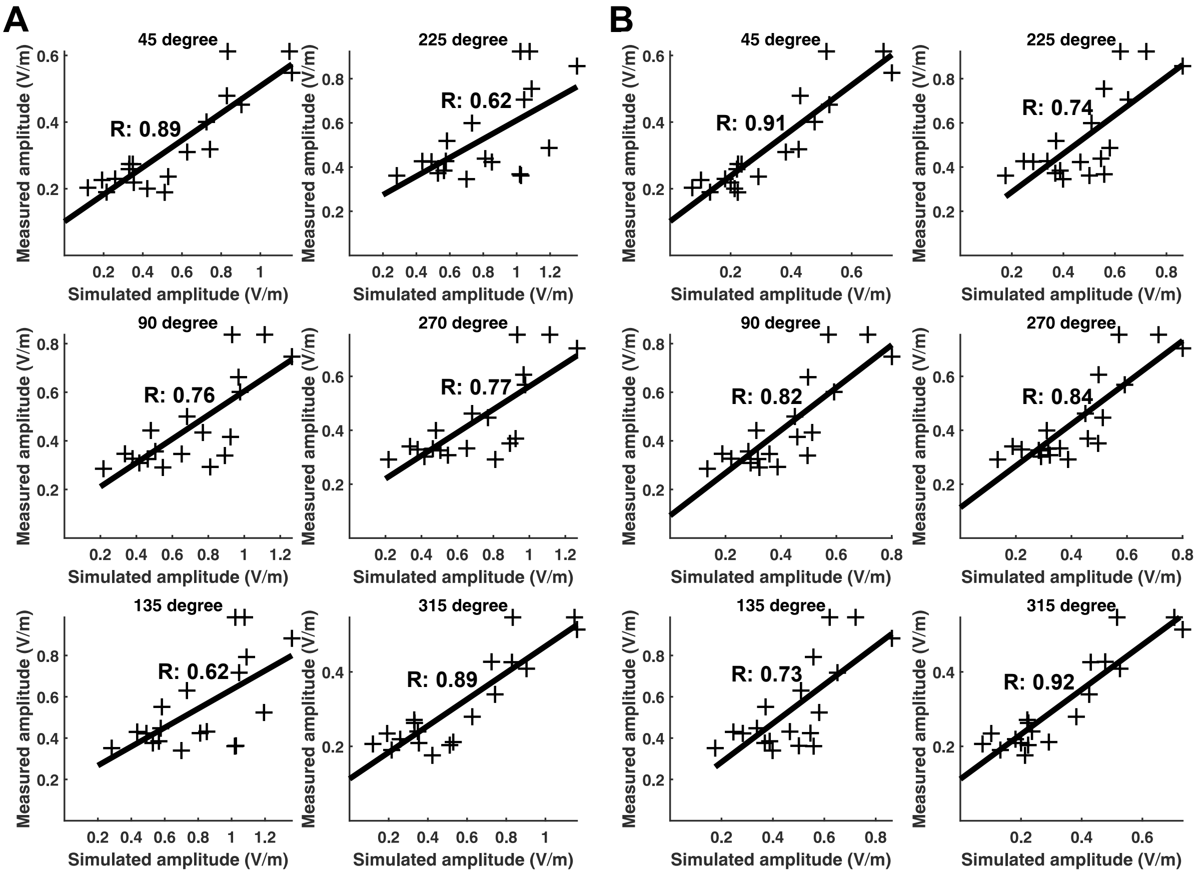


**Supplementary Figure. 11.** Correlation between the measured amplitude and either A) the simulated amplitude obtained when applying the initial conductivity or B) the simulated amplitude calculated with the optimal conductivity for monkey 2.
